# A mutant RNA polymerase activates the general stress response, enabling *E. coli* adaptation to late prolonged stationary phase

**DOI:** 10.1101/2020.01.27.922484

**Authors:** Pabitra Nandy, Savita Chib, Aswin Seshasayee

## Abstract

*Escherichia coli* populations undergo repeated replacement of parental genotypes with fitter variants deep in stationary phase. We isolated one such variant, which emerged after three weeks o of maintaining an *E. coli* K12 population in stationary phase. This variant displayed a small colony phenotype, slow growth and was able to outcompete its ancestor over a narrow time window in stationary phase. The variant also shows tolerance to beta-lactam antibiotics, though not previously exposed to the antibiotic. We show that an RpoC (A494V) mutation confers the slow growth and small colony phenotype to this variant. The ability of this mutation to confer a growth advantage in stationary phase depends on the availability of the stationary phase sigma factor σ^S^. The RpoC (A494V) mutation up-regulates the σ^S^ regulon. As shown over 20 years ago, early in prolonged stationary phase, σ^S^ attenuation, but not complete loss of activity, confers a fitness advantage. Our study shows that later mutations enhance σ^S^ activity, either by mutating the gene for σ^S^ directly, or via mutations such as RpoC (A494V). The balance between the activities of the housekeeping major sigma factor and σ^S^ sets up a trade-off between growth and stress tolerance, which is tuned repeatedly during prolonged stationary phase.

**Importance:** An important general mechanism of bacterial adaptation to its environment involves adjusting the balance between growing fast, and tolerating stresses. One paradigm where this plays out is in prolonged stationary phase: early studies showed that attenuation, but not complete elimination, of the general stress response enables early adaptation of the bacterium *E. coli* to the conditions established about 10 days into stationary phase. We show here that this balance is not static and that it is tilted back in favour of the general stress response about two weeks later. This can be established by direct mutations in the master regulator of the general stress response, or by mutations in the core RNA polymerase enzyme itself. These conditions can support the development of antibiotic tolerance though the bacterium is not exposed to the antibiotic. Further exploration of the growth-stress balance over the course of stationary phase will necessarily require a deeper understanding of the events in the extracellular milieu.

## Introduction

Under standard laboratory conditions, well-aerated cultures of *Escherichia coli* exhibit three distinct phases of growth within 24 hours of inoculation: the initial slow growing *lag* phase, followed by a period of *exponential* growth and finally the *stationary* phase during which no appreciable change in cell number is observed. Most of the cells in culture lose viability within 2-5 days after reaching stationary phase [(1), (2)]. The small surviving subpopulation can remain viable for at least 60 months in a phase known as “prolonged stationary phase” or “long-term stationary phase” (1, 3, 4). Prolonged stationary phase might represent a laboratory approximation of different natural habitats populated by bacteria, many of which are nutrient limited [(2)].

Although the total number of viable cells does not appreciably change in this phase, prolonged stationary phase is dynamic and is characterized by repeated replacement of parental genotypes (and phenotypes) by a different set of genetic variants better adapted to the ambient environment [(2)]. Populations from aged cultures outcompete younger cultures in prolonged stationary phase, a phenotype known as Growth Advantage in Stationary Phase (GASP) [(4)]. In this phase, both genetic and phenotypic diversity increase over time (2, 5).

Mutations in different global regulators of gene expression confer GASP on their host cells. The most prominent among these are in the sigma factor *σ^S^*, the master activator of the general stress response. An elongated and attenuated variant of *σ^S^*, named *rpoS819,* was the first gene to be implicated in GASP [(4)]. Other alleles conferring GASP include mutations in Lrp, a global regulator controlling amino acid metabolism [(6)]; activation of the *bgl* operon [(7)] by mechanisms that attenuate H-NS, the global regulator responsible for transcriptional repression of horizontally acquired genes [(8)]; and mutations in CpdA, the cyclic-AMP phosphodiesterase [(9)], which might influence hierarchy of carbon source utilization. Several mutations in the core RNA polymerase subunits have been observed in deep sequencing based studies of the genotype space explored in prolonged stationary phase [(5), (10), (11)]. Previous research has described the emergence of diverse colony morphologies in prolonged stationary phase [(3)]. Consistent with these observations, we isolated a variant characterized by its small colony size growing alongside cells forming ‘normal’ sized colonies, after maintaining *E. coli* in stationary phase for over 3 weeks.

Slow growing mutants of various bacteria can withstand a variety of stresses targeting growth-associated processes. For example, slow growing *E. coli* variants deficient in the electron transport chain can tolerate aminoglycosides whose transport into the cells is an active process (12). A certain class of slow growing bacteria, called “Small Colony Variants (SCV)” emerge in response to stress in the environment [(13)], and have been discovered in pathogenic contexts. These variants arise from metabolic deficiencies, particularly in thymidine biosynthesis or in the electron transport chain. Some of these variants are known to evade host defence systems and the action of various antibiotics, while forming persister populations responsible for recalcitrant diseases [(14)].

In this study we describe the phenotypic characteristics and genetic underpinnings of a slow growing, small colony forming variant of *Escherichia coli* isolated three weeks into stationary phase in nutrient rich medium (LB), and show that a mutation in the core RNA polymerase can confer GASP to its bearer by activating the *σ^S^* regulon.

## Results

### Small colony phenotype evolved from ZK819 in prolonged stationary phase

In a previous study [(5)], we had propagated 4 replicate populations of *Escherichia coli* ZK819 in rich media (LB) for 30 days without the addition of external nutrients after inoculation. ZK819 is the pioneer GASP strain of *E. coli*, carrying *rpoS819,* an elongated and attenuated variant of the sigma factor for the general stress response *σ^S^*. In this experiment, colonies with different morphologies emerged over time: mucoid colonies, followed by fragmented colonies and small sized colonies. Variants with small colony (SC) phenotype first appeared in two replicate lines on day 22 and remained for as long as the cultures were examined (Figure 1) [(5)]. We isolated a SC from these long-term stationary phase experiments for further study, and describe its characteristics here.

**Figure 1:**
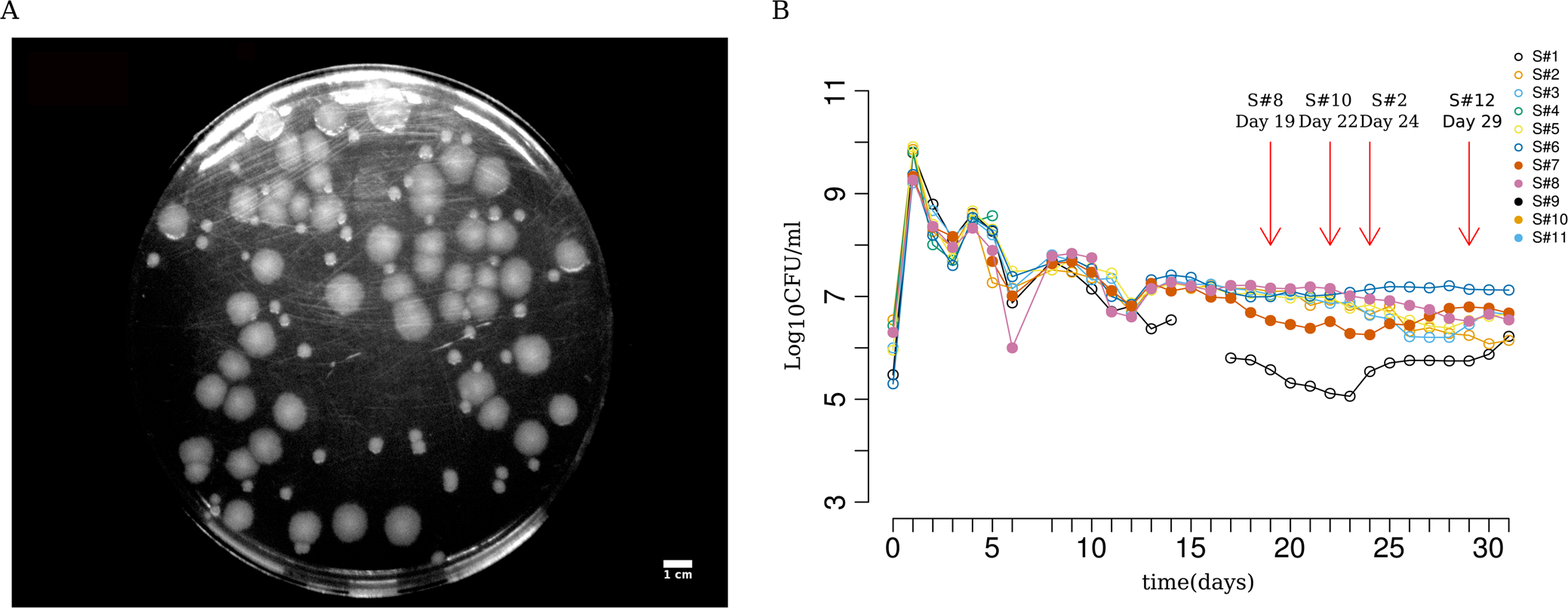
Evolving a GASP-enabled strain (ZK819) in prolonged stationary phase and the emergence of small colony forms. (A) Representative picture of small colonies as seen on LB agar plate spread with 29 day old culture of ZK819. (B) Daily colony count of 12 independent replicates of *E. coli* ZK819 for 30 days in prolonged stationary phase. First recorded emergence of small colonies in different replicate lines are marked in red arrows.

We briefly note here that the appearance of the SC phenotype is repeatable, and we observed its appearance, after 19-22 days, in four out of 11 additional prolonged stationary phase runs. These were not studied in greater detail here.

### SC exhibits slow growth and increased tolerance to beta-lactam antibiotics as compared to ancestor and co-existing strains

We measured the growth of SC, a co-existing variant with ancestral colony size (LC) and the parental ZK819 in LB (Figure 2A). SC exhibited slow growth, both in terms of maximum growth rate (Figure 2C) and lag times as compared to ZK819 and LC (Figure 2D). All three strains showed similar stationary phase yields in rich media. In minimal media supplemented with glucose as sole carbon source, SC retained its slow growth, additionally showing lower yield compared to other strains (Figure 2B).

**Figure 2:**
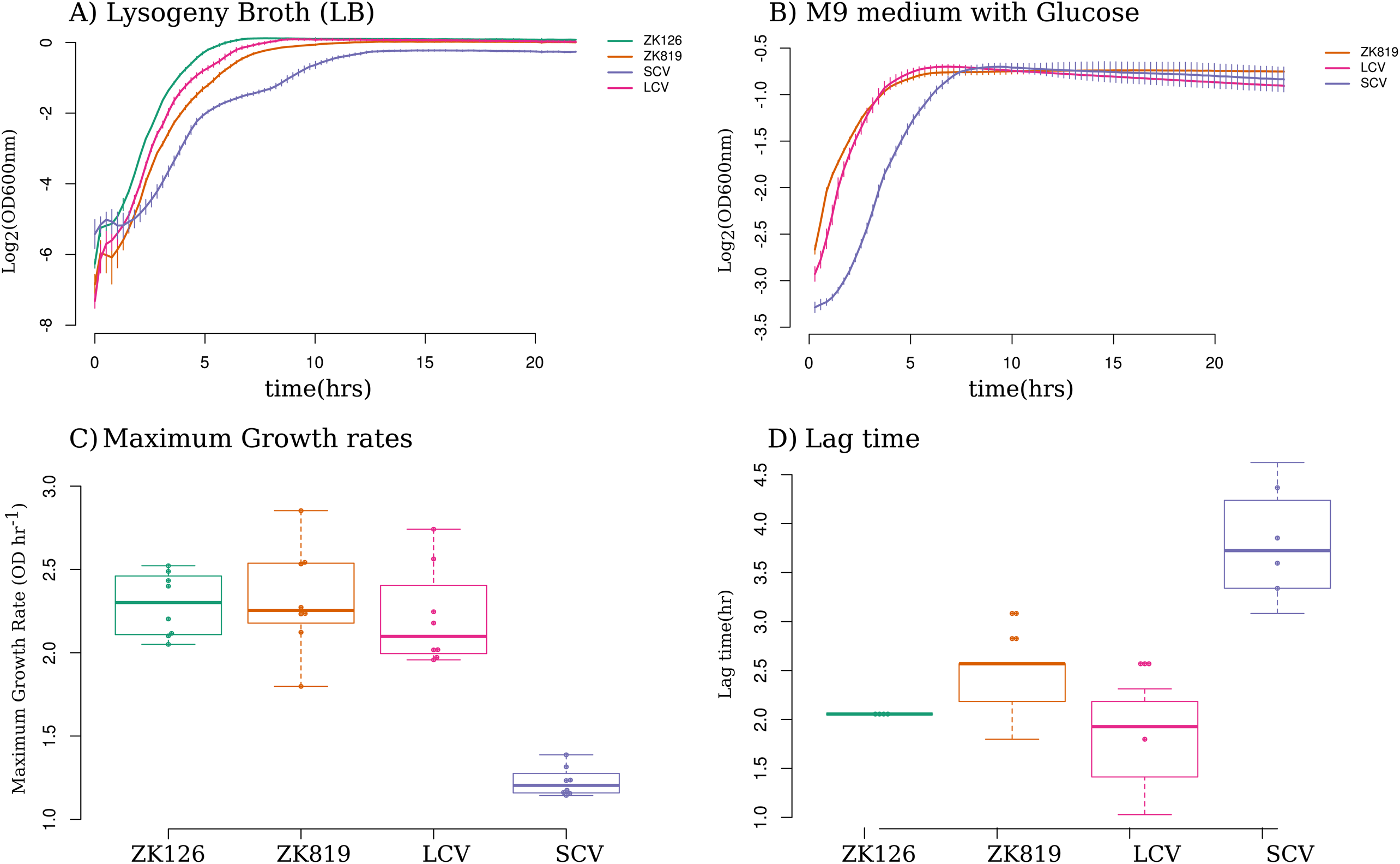
Growth characteristics of SC. Growth curves of SC compared to different parent strains in (A) rich (LB) and (B) minimal (M9+glucose) media. (C) Distribution of maximum growth rates of SCV, LCV and ancestor strains. Error bars indicate standard deviation (N=8). (D) Distribution of lag time for SCV, LCV and ancestor strains. Error bars indicate standard deviation(N=8).

Small colony variant populations isolated in other bacteria - often in pathogenic contexts - show increased tolerance to antibiotics [(15), (16)]. We tested if this would apply to the SC isolated here by comparing the minimum inhibitory concentration (MIC) of three different antibiotics - ampicillin, kanamycin and ciprofloxacin - against SC, LC and ZK819. SC showed an increased resistance to ampicillin and the analogous carboxy-penicillin antibiotic carbenicillin, but not to the other two antibiotics (Figure 3, *Supplementary Figure S1*).

**Figure 3:**
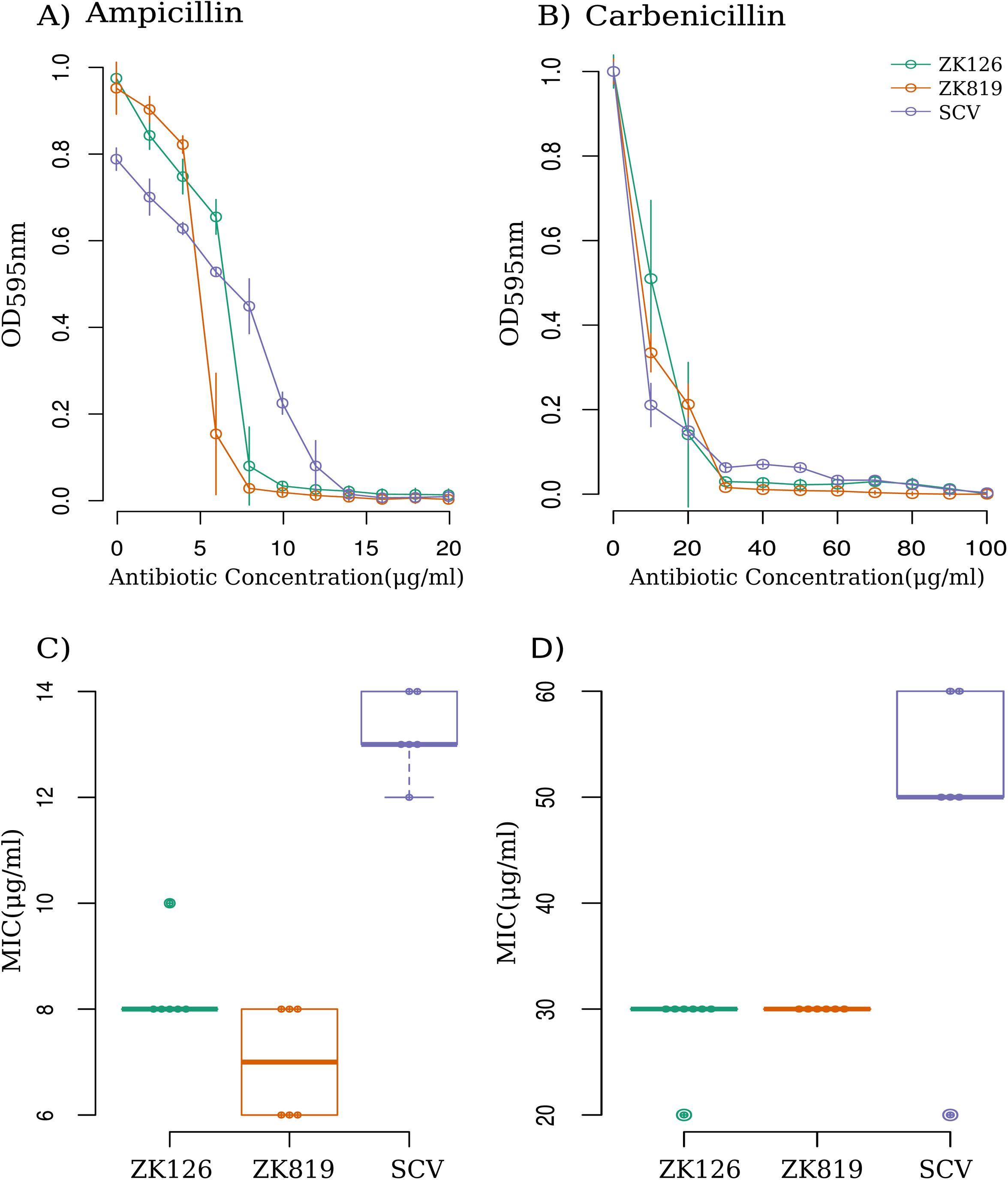
SC exhibits tolerance against beta lactam antibiotics. Kill-dynamics of (A) Ampicillin and (B) Carbenicillin against SC and ancestor strains. Distribution of MICs of evolved mutant (SCV), parent (ZK819) and ancestor (ZK126) strains against (C) Ampicillin and (D) Carbenicillin (N=6).

We observed that ampicillin clears more SC as compared to wild type when present in lower concentrations. At higher concentrations of the antibiotic this trend is reversed. This type of concentration-dependent variable response requires different subpopulations to have variable affinity to the antibiotic and is known as “heteroresistance” [(17)].

### SC shows GASP phenotype in a limited time window in prolonged stationary phase

Growth Advantage in Stationary Phase (GASP) is defined as the ability of a strain to outcompete its ancestor in prolonged stationary phase [(4)]. To examine the ability of SC to show GASP, we competed SC against its ancestor ZK819 using mixed culture competition experiments [(4)] in fresh LB and in media spent for different periods with the ancestral strain ZK819. In these experiments, the competing strains were inoculated in three different ratios: 1:1000, 1:1 and 1000:1.

We found that SC does not show any competitive advantage against ZK819 in freshly autoclaved and 3 day-old spent media. However, it outcompetes ZK819 in 10 day-old media. Starting from 1000-fold minority, SC overtakes ZK819 after three days only in the 10 day-old spent medium (Figure 4). Though it is not unreasonable to expect considerable batch-to-batch variation in spent media across trials, the above result was consistently found across three replicates. In 20 day and 30 day-old media however, the competitive advantage of SC was lost, indicating that SC can exhibit GASP only in a narrow time window in prolonged stationary phase (*Supplementary Figure S2*). This is consistent with the idea that the continuous wax and wane of different genetic variants is a reflection of the ever-changing biochemical properties of the prolonged stationary phase medium [(3)].

**Figure 4:**
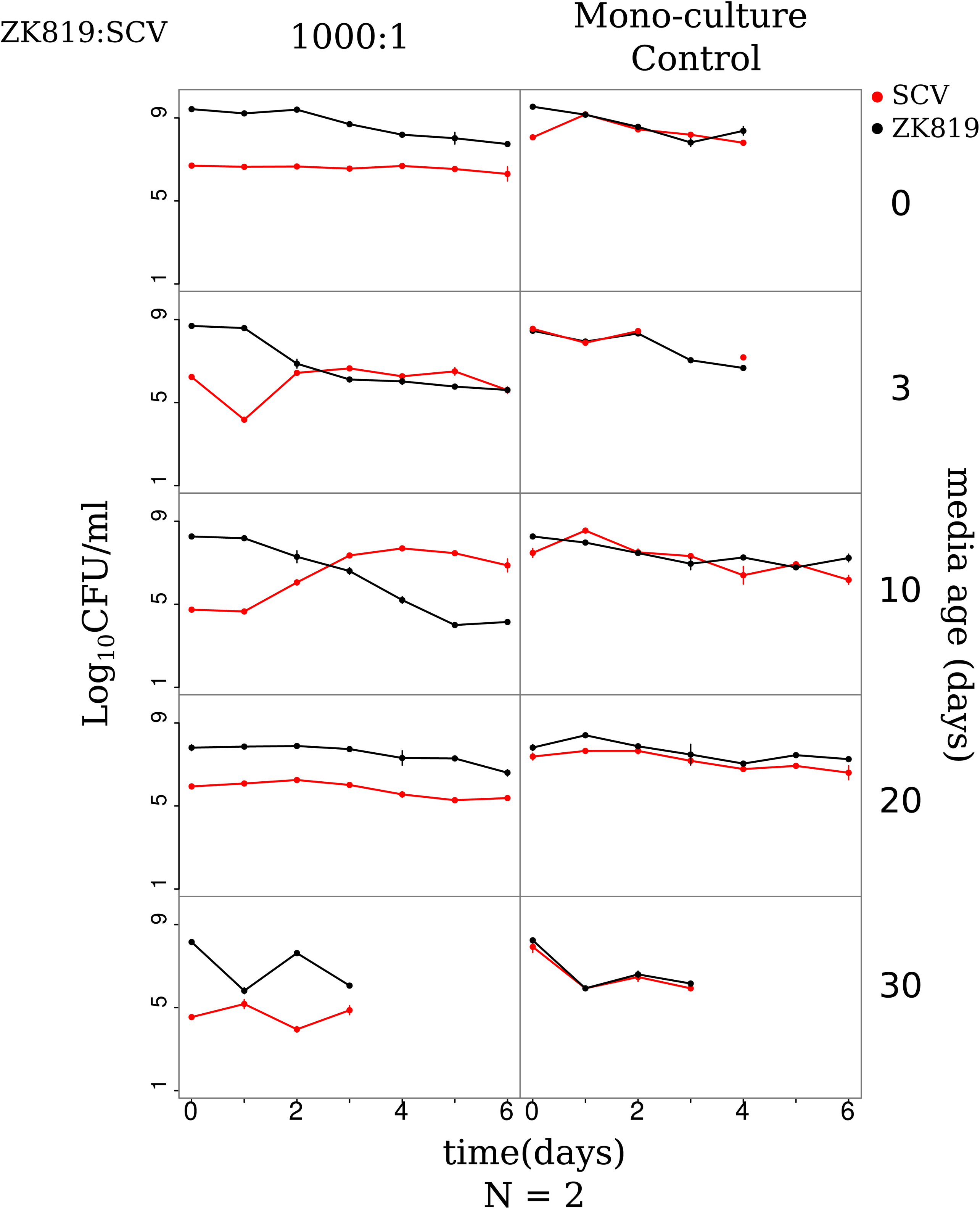
SC exhibits competitive advantage against its ancestor within a specific time window in prolonged stationary phase. Competition dynamics between ZK819(ancestor) and SC in different aged LB. Left panel shows competition experiments where ZK819 are inoculated in 1000 times higher frequency compared to SC. Right panels shows competing strains grown individually in the same aged media as controls. Age of media used for competition is shown on right. Competition dynamics with other starting frequencies are supplied in Supplementary Figure S2.

In light of the increasing genetic diversity of the population and the late emergence of SC, it is more reasonable to expect SC to compete with the coexisting LC than with the ancestral ZK819. We therefore investigated the competitive advantage of SC against LC in 10 day-old LB, and found that SC, starting from a 1000-fold minority, overtook LC after two days (*Supplementary Figure S3*). Therefore, SC shows the ability to outcompete both ZK819 and LC in prolonged stationary phase in conditions representing a narrow time-window.

### rpoC^A494V^ is a unique mutation in SC that causes small colony size, irrespective of background

To identify and characterize the genetic basis of the small colony and the GASP phenotype of SC, we performed whole-genome sequencing of SC, LC and ZK819. We found several mutations common to SC and LC, and identified two loci at which the two diverged. The first was at *rpoS*, encoding *σ^S^*. Whereas ZK819 carried the attenuated *rpoS819* allele, SC carried what we refer to as *rpoS92. rpoS819* is defined by a 46 bp duplication at the 3’ end of the gene. This duplication results in a longer protein with reduced activity, which may in part be due to its reduced expression at the protein level. *rpoS92* carries a reduplication of the 46 bp region. This introduces a STOP codon prematurely for *rpoS819*, resulting in a *σS* protein variant of length intermediate between the wild type and *rpoS819*. *rpoS92* partially restores the activity of *σ^S^* that had been lost in *rpoS819* [(9)]. LC, in contrast to both SC and ZK819, carries the wild type *rpoS* ORF.

The second locus at which these three strains diverged was *rpoC*, coding for the β’ subunit of the core RNA polymerase. Whereas ZK819 and LC carry the wild type *rpoC* locus, SC has a G:A transition, resulting in the replacement of the alanine at residue 494 by a valine. Mutations in the core RNA polymerase have been found to confer adaptive advantages to *E. coli* under different conditions, and have been found in at least two whole genome sequencing studies of variants that emerge in prolonged stationary phase [(5), (10)]. We also identified a unique synonymous mutation (H366H) in LC in the gene *icd* coding for isocitrate dehydrogenase, which we have not characterized in this study.

To test the phenotypic effect of *rpoC^A494V^* mutation, we introduced it in different relevant genetic backgrounds using phage mediated transduction from a *thiC39::tn10* marker (CAG18500), as described in a previous study [(18)]. The genetic backgrounds in which *rpoC^A494V^* was introduced were selected based on their *σ^S^* status and their relevance to the strains used or isolated in this study; these were LC (evolved strain, *rpoS^WT^*), ZK819 (ancestor, *rpoS819*), ZK126 (*rpoS^WT^*, the parent of ZK819) and *ΔrpoS* (ZK819::*ΔrpoS*). In addition, the wild type *rpoC* allele was transduced into SC reverting the *rpoC^A494V^* mutation to the wildtype. The presence of the *rpoC^A494V^* mutation was confirmed by sanger sequencing with relevant primers (*Supplementary Table S1*). Single-end whole genome sequencing were performed on all transductant strains to check for the presence of any additional indels, copy number variations and point mutations that may have been inadvertently introduced during transduction and found none apart from those introduced.

We observed that *rpoC^A494V^* bearing transductants in all backgrounds, including *ΔrpoS,* produced small colonies on LB agar plates and exhibited slow growth, characteristic of SC in liquid culture (Figure 5 A, B). Conversely, replacing the mutant *rpoC^A494V^* allele in SC background with the wild type *rpoC* allele ameliorated slow growth, thus reverting the small colony phenotype to ancestral colony size. These results indicate that the *rpoC^A494V^* mutation is sufficient to cause the small colony phenotype to its bearer, independent of the status of *σ^S^*.

**Figure 5:**
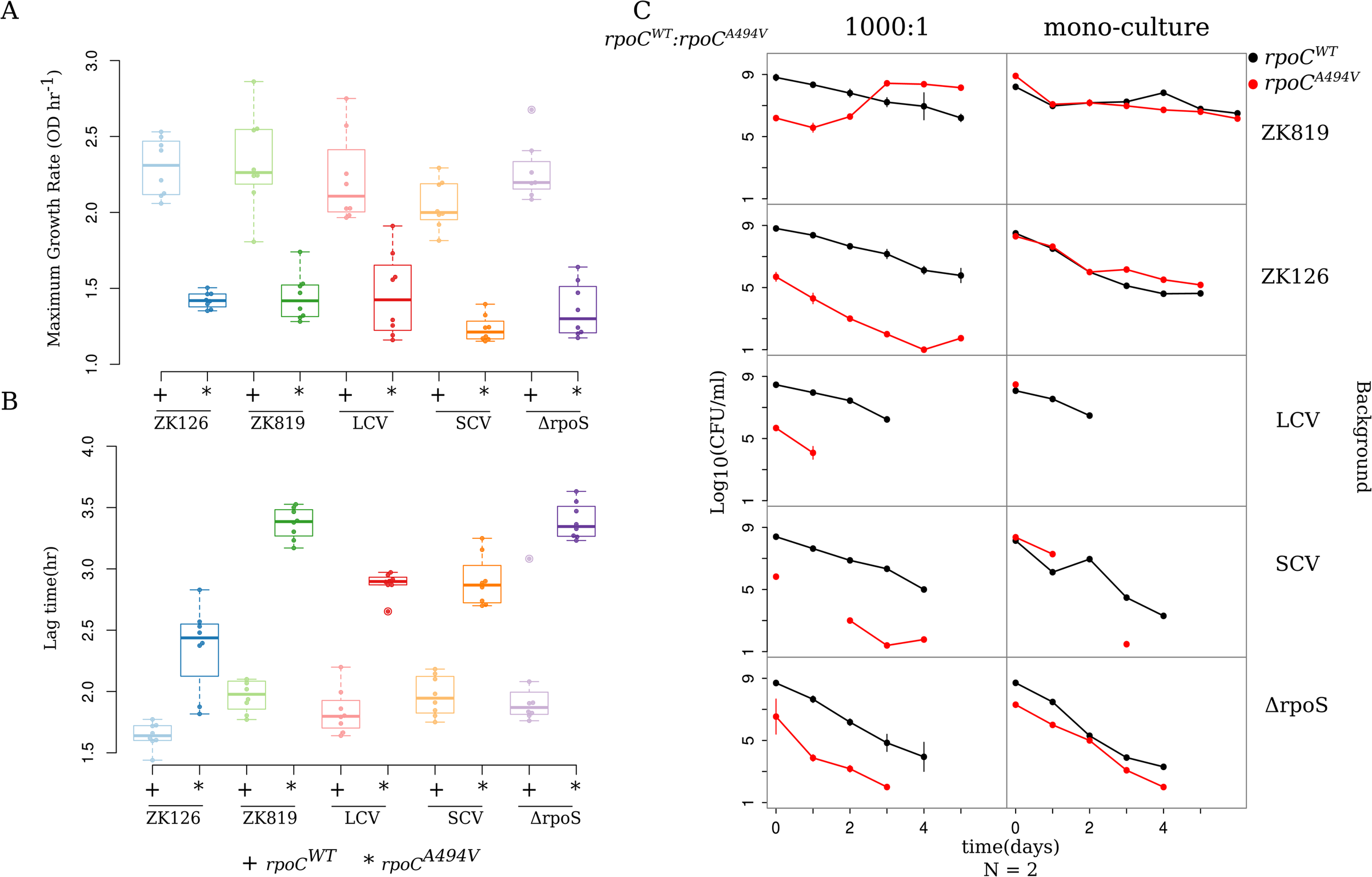
rpoCA494V is necessary and sufficient to cause a slowdown in growth and confers GASP in a background with attenuated *σ^S^* activity. (A) Comparison of Maximum growth rates of strains carrying RpoC^WT^ (marked +) and rpoC^A494V^ (marked *) in different backgrounds. Maximum growth rates were calculated from growth data as described in Materials and Methods. (B) Comparison of lag times of strains carrying RpoC^WT^ (+) and RpoC^A494V^ (*) in different backgrounds. Lag times were calculated from growth curve data as in Materials and Methods. (C) Competition dynamics between strains carrying RpoC^WT^ and RpoC^A494V^ in different backgrounds. Left panel shows scenarios where RpoC^WT^ strain is inoculated in 1000 times higher frequency compared to RpoC^A494V^ strain. Right panels shows competing strains grown individually in the same age media as controls. Data shown only for competitions performed in 10 day old spent media. Other media conditions and starting frequencies are provided in Supplementary Figures 4-8.

### rpoC^A494V^ mutant might cause inefficient transcription

The ability of *rpoC^A494V^* to slow growth is independent of the status of *σ^S^*. This being a mutation in the core RNA polymerase, we suspected that it would have a broad impact on global gene expression levels.

To test whether the mutation causes any alteration to transcription, we tested induction kinetics of the chromosomal *ara* operon in *rpoC* mutant and wild type strains in ZK819 background. Note that the *ara* operon comes under the control of the housekeeping sigma factor *σ^D^* and not *σ^S^*. We measured the induction of the *araB* gene over time following the addition of arabinose to an exponentially growing culture by performing RT-PCR using primers amplifying a 134 bp region at the 5’ end of the gene (*Supplementary Table S1*). To limit any population-level heterogeneity in *ara* induction [(19)], we used nearly 100-fold excess of arabinose for induction.

In *rpoC^WT^*, *araB* levels increase steeply (ΔΔFCinduced-uninduced = −6.05) within 50 minutes of adding arabinose. In *rpoC^A494V^*, increase in mRNA is muted (ΔΔFCinduced-uninduced = ranging from −0.74 to −4.54), more variable across biological replicates, and delayed, occurring after 110 minutes post induction (Figure 6A). This delayed and repressed response to arabinose induction in *rpoC^A494V^* suggests that the mutation reduces the activity of RNA-pol. This is unlikely to be an effect of a possible reduced arabinose transporter expression in the mutant: RNA-seq data (see below), showed no significant difference in the basal expression level of *araE* between the wild type and mutant *rpoC* strains. Thus slow transcription might underpin the ability of *rpoC^A494V^* to cause slow growth.

**Figure 6:**
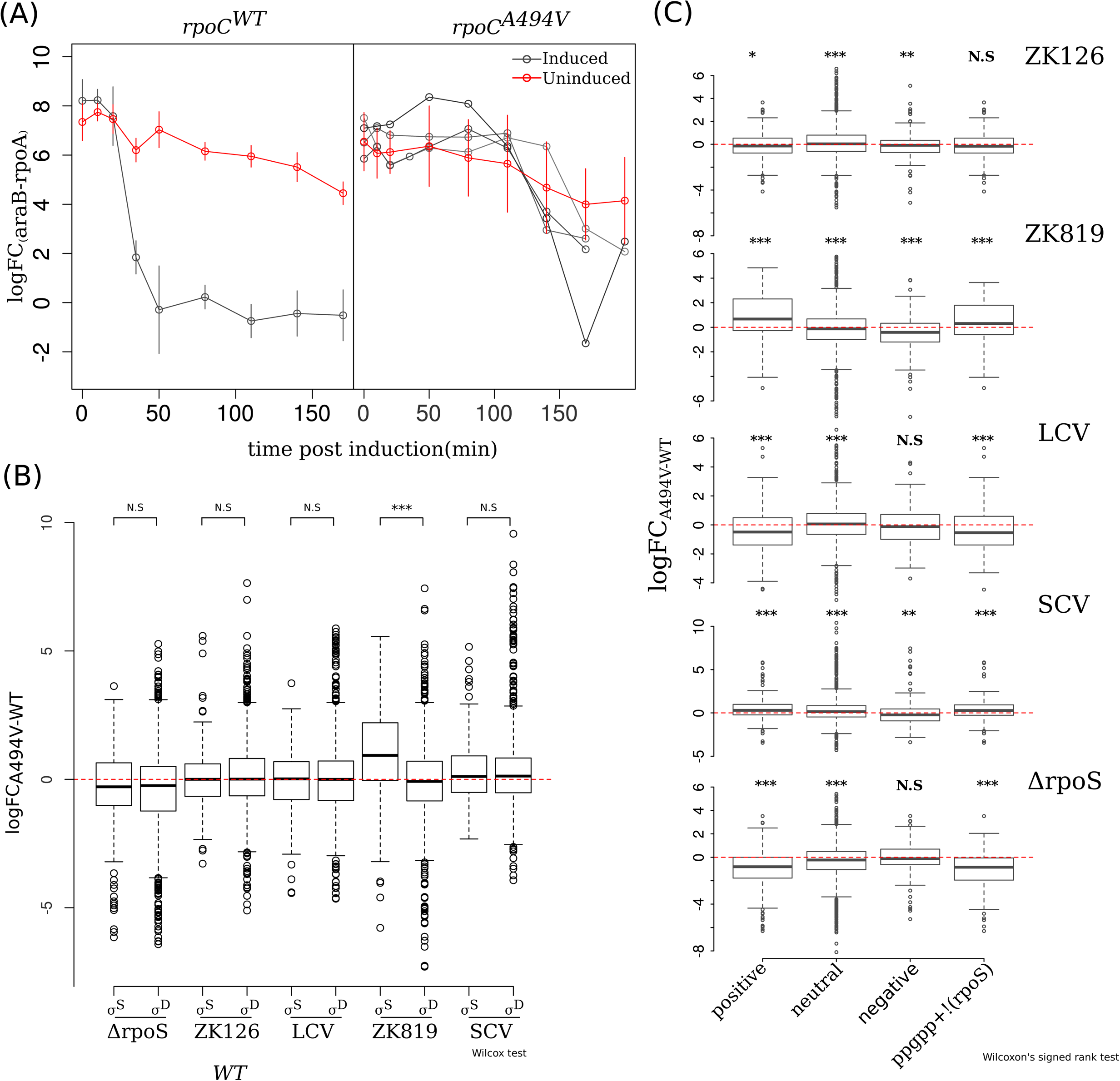
RpoC^A494V^ mutation affects global gene expression states. (A) Semi-quantitative estimation of *araB* expression by RT-PCR in RpoC^WT^ (left) and RpoC^A494V^ (right) in the ZK819 background, post induction with 100mM of L-Arabinose. Error bars indicate Standard deviation (N=4). Individual Replicates have been shown separately in RpoC^A494V^ strain to emphasize the biological variation observed in the induction pattern in mutant strain compared to wild type. Ct values from *araB* mRNA was normalized with that of constitutively expressing *rpoA* gene. *araB* measurements from uninduced cultures as controls are shown in red. (B) Differential expression of σ^S^ and σ^D^ target genes between RpoC^A494V^ and RpoC^WT^ in different backgrounds. Wilcoxon’s test was used to test significant of the difference in median expression values. p-value scale ***<0.001, **<0.01, *<0.05. (C) Differential expression of positive, negative and neutrally regulated targets of ppGpp between RpoC^A494V^ and rpoC^WT^ in different backgrounds (noted in top right of each figure). To control for the behaviour of a large number of genes controlled by both σ^S^ and ppGpp, differential expression of genes which are upregulated by ppGpp but are not σ^S^ targets were taken also included.

### Competitive advantage conferred by rpoC^A494V^ mutation depends on the status of σS

Having established that the *rpoC^A494V^* mutation can cause the small colony phenotype independently of the status of *σ^S^*, we interrogated the ability of this mutation to confer GASP to its bearer. To do this, we performed mixed culture competition experiments of *rpoC^A494V^* strains against *rpoC^WT^* strains in all relevant genetic backgrounds in both fresh and spent media.

We found that *rpoC^A494V^* confers GASP on its bearer only in the ZK819 background, carrying the attenuated *rpoS819* allele, and not others (Figure 5C, *Supplementary Figures S4-8*). In 10 day-old spent media, ZK819+*rpoC^A494V^* overtook ZK819+*rpoC^WT^* from a 1000 folds lower starting frequency after two days. However, SC with *rpoS92* and *rpoC^A494V^* did not outcompete SC with *rpoS92* and *rpoC^WT^*. This suggests, under the assumption that these mutations emerged under a regime of natural selection, that the *rpoC^A494V^* mutant might have arisen in the *rpoS819* background, with *rpoS92* emerging later. We also found that in the ZK819*ΔrpoS* background, both *rpoC^A494V^* and *rpoC^WT^* exhibited precipitous drops in cell count with no viable colonies remaining after three days of growth, highlighting the importance of *rpoS* in survival and growth in nutrient depleted/altered conditions (*Supplementary Figure S4*).

These suggest that *rpoC^A494V^* confers GASP only when *σ^S^* is severely attenuated, as seen in *rpoS819*. However, its ability to confer GASP does require some level of functional *σ^S^*, bringing about the suggestion that it acts through *σ^S^*.

### *rpoC^A494V^* induces expression of the *σ^S^* regulon depending on the *rpoS* genetic background

Towards understanding the mechanisms by which *rpoC^A494V^* causes small colony and the GASP phenotype, we performed RNA-seq based transcriptome experiments comparing *rpoC^A494V^* and *rpoC^WT^* in different genetic backgrounds differing in their *σ^S^* activities (ZK126, ZK819, SC, LC and *ΔrpoS*). We identified differentially expressed genes that showed a log2-fold change of over 1.5 at P-value of less than 0.05.

The number of differentially expressed genes across these five backgrounds ranges from 407 to 654. However, the number of genes differentially expressed irrespective of the genetic background was only 18. This shows that the effect of *rpoC^A494V^* on the transcriptome is dependent on the background expression state established at least in part by *σ^S^* activity.

Because the ability of *rpoC^A494V^* to confer GASP - as well as alter gene expression states - seemed to depend on the status of *σ^S^*, we tested the effect of the mutation on the expression levels of *σ^S^* targets. Towards this, we compared the fold changes - between *rpoC^A494V^* and *rpoC^WT^* - of targets of *σ^S^* with those of a set of targets of the housekeeping sigma factor *σ^D^*. In a second analysis, we tested for the overlap between *σ^S^* and σ^D^ target genes that are differentially expressed in *rpoC^A494V^* over *rpoC^WT^*. When we looked at all genes, *σ^S^* targets showed higher positive fold changes in *rpoC^A494V^* when compared with *rpoC^WT^*, only in the ZK819 background carrying the attenuated and elongated *rpoS819* allele (Figure 6B). Consistent with this, sets of differentially expressed gene sets between *rpoC^A494V^* and *rpoC^WT^* were significantly enriched for *σ^S^* targets in ZK819 (*rpoS819*) background, but not the other backgrounds (*Supplementary Table S2-S6*).

A previous study had identified *rpoC^A494V^* as a mutation that enables RNAP to bypass the requirement of the small molecule ppGpp in transcribing a ppGpp dependent, *σ^54^* promoter [(18)]. Following from this work, we analysed the expression patterns of genes known to respond to ppGpp [(20)] in *rpoC^A494V^* compared to *rpoC^WT^*. We observed that positively regulated targets of ppGpp are upregulated and negatively regulated targets down regulated in *rpoC^A494V^* compared to *rpoC^WT^* in the ZK819 and SC backgrounds, which harbor elongated *rpoS* alleles with less than wild type *σ^S^* activity (Figure 6C). This applies for genes that are known to respond to ppGpp, but not known to be regulated by *σ^S^* as well. While this might indicate that this effect may not be mediated by *σ^S^*, we observed an opposite pattern in *rpoC^A494V^* compared to *rpoC^WT^* in the *ΔrpoS* background (Figure 6C). We did not observe any significant effect on ppGpp targets in the *rpoS^WT^* background.

*rpoC^A494V^* does not seem to induce the *σ^S^* regulon by increasing the expression levels of *σ^S^*, on the basis of our observation that there is little differential expression of the *rpoS* gene in the RNA-seq data, nor is there any increase in *σ^S^* protein levels as measured by Western blotting. In fact, Western blots indicate a decrease in the levels of *σ^S^* in *rpoC^A494V^* (Figure 7). This suggests that *rpoC^A494V^* may activate the *σ^S^* regulon by affecting σ-factor competition for the core RNA polymerase. There is evidence [(21)] that ppGpp reduces the ability of *σ^D^* to compete with at least two alternative sigma factors including *σ^S^*. Whether the induction of the *σ^S^* regulon by *rpoC^A494V^* operates through this mechanism, and whether this might even require the extended C-terminal tail that *rpoS819* and *rpoS92* alleles add to the wild type *rpoS*, remains to be tested.

**Figure 7:**
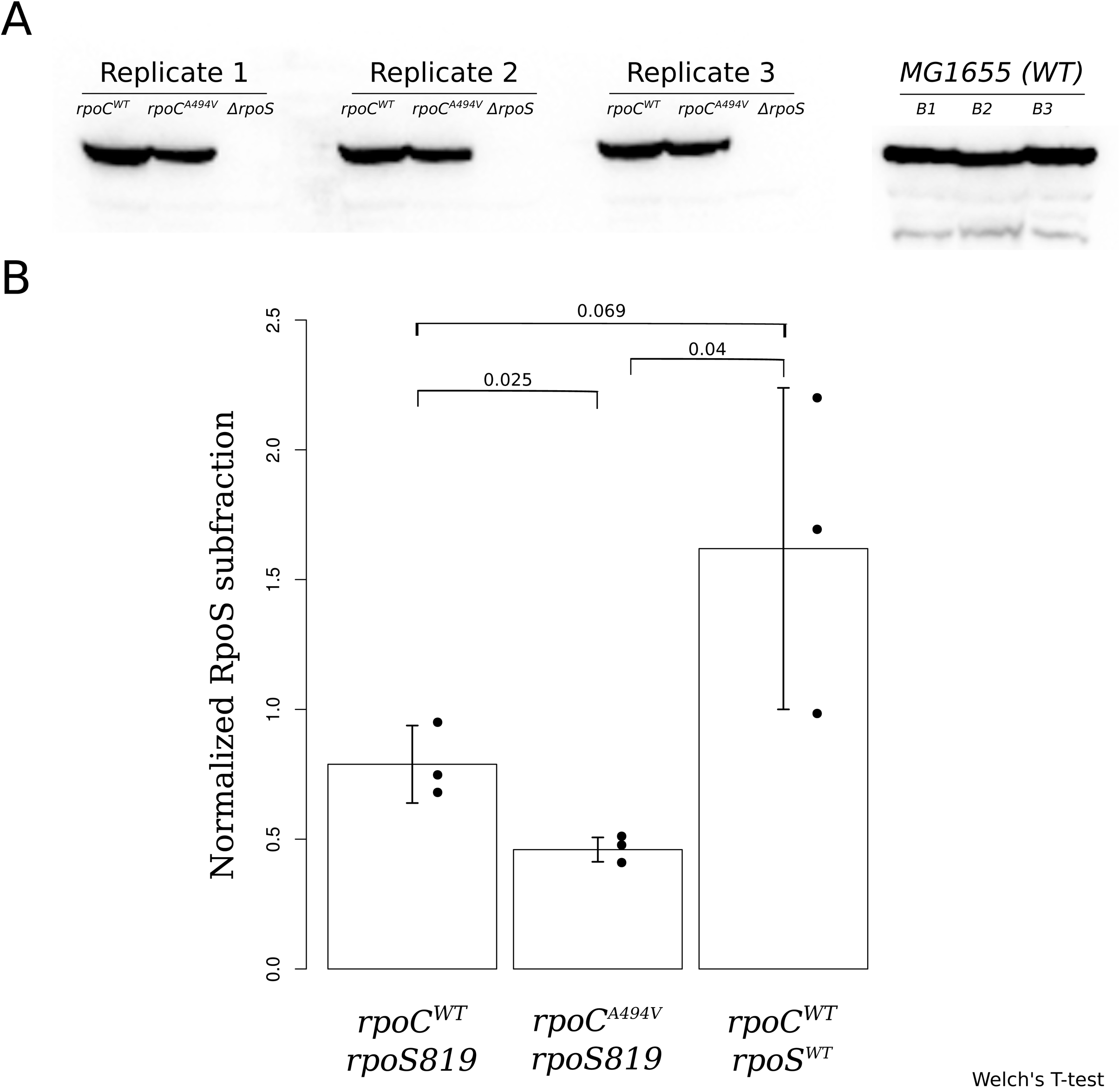
*σ^S^* protein level in RpoC^WT^ and RpoC^A494V^ strains in ZK819 background. (A) Nitrocellulose blot containing *σ^S^* protein in RpoC^WT^ and RpoC^A494V^ strain the *rpoS819* background. *ΔrpoS* strain, with the deletion introduced into the ZK819, is shown as control. On the right, *σ^S^* protein derived from wild-type MG1655 strain containing *wt rpoS* allele is shown as positive control. A part of the gel was stained with coomassie blue and total protein intensities from each lane was quantified which was then used for normalizing the rpoS protein intensity (Supplementary Figure S9). (B) Normalized intensity of *σ^S^* protein was quantified for each strain and average value is shown. Error bars represent standard deviation over three replicates. Statistical significance of difference of intensity between each pair of strains were calculated by Welch’s T-test and p-values are shown.

In summary, though *rpoC^A494V^* increases the expression of the *σ^S^* regulon in backgrounds with low *σ^S^* activity, this effect requires the presence and expression of *σ^S^*. The manifestation of GASP in the regime in which *rpoC^A494V^* displays a selective advantage might require higher *σ^S^* activity, in contrast to earlier time points in which attenuated variants of *σ^S^* such as *rpoS819* are favoured. In backgrounds with elongated *rpoS* alleles the effect of the *rpoC^A494V^* allele on the transcriptome shows similarities to that of higher intracellular ppGpp. This effect also seems to be mediated by *σ^S^* as in the *ΔrpoS* background, the effect of *rpoC^A494V^* mutation flips and resembles that of lower intracellular ppGpp.

## Discussion

This work highlights the global effect of a single nucleotide substitution in the β’ subunit of bacterial RNA polymerase emerging a few weeks into prolonged stationary phase. We show that in this phase, slow growth is advantageous within a certain time window. Our experiments demonstrate the *rpoC^A494V^* mutation is necessary and sufficient to confer slow growth to the bacteria. However, to show GASP phenotype, attenuated *rpoS819* allele is additionally required.

Genetic and phenotypic diversification of an initially isogenic inoculum is a characteristic of prolonged stationary phase. Constantly changing physico-chemical condition of the media results in different sub-populations arising over time. This is orchestrated by different allelic forms of *σ^S^* that sequentially emerge with time. Within the first 10 days, the elongated and attenuated *rpoS819* allele emerges giving rise to the first GASP population [(4)]. A constant competition for occupying the RNA polymerase holoenzyme exists between the housekeeping sigma factor *σ^D^* driving growth and the stationary phase sigma factor *σ^S^* involved in dormancy and maintenance [(22), (23)]. There is always a trade-off between growth and stress response, which is determined by this competition between the two sigma factors. The expression and activity of *σ^S^* is tightly regulated at multiple levels, and adjusting this switch helps the bacterial cell modulate global gene expression states according to environmental cues. [(24, 25)]. Additionally, the gene for *σ^S^* is highly mutable, thus enabling genetic alterations of the growth-stress tolerance trade-off (26).

The attenuated *rpoS819* allele alters this growth-stress response tradeoff in favour of growth. In our experiments we found that the *rpoS819* allele, when evolved in LB media, suffers another re-duplication at C-terminal end around 19 −22 days post inoculation, giving rise to *rpoS92* allele which has activity closer to that of the wildtype *rpoS* allele [(9)]. The *rpoC^A494V^* mutation also enhances the activity of *σ^S^*. This suggests that stress response takes some precedence in later stages of stationary phase. GASP outcomes depend on initial media conditions [(27)], and this can interplay with the balance between stress tolerance and growth.

Mutations in the *rpoABC* operon results in a wide variety of pleiotropic phenotypes, by affecting different steps of transcription initiation process [(28)]. In a study by Mukherjee *et. al* mutations mapped nearby to locus of our mutation of interest on the β’ subunit are found to confer resistance to antibacterial micropeptide Microcin J25 [(29)]. This region lies in the secondary channel of RNA polymerase in the shape of a “hollow cylinder”. Our mutation of interest *RpoC^A494V^* could alter the local architecture of the secondary channel, reducing the rate of influx of ribonucleotides to be assembled to a nascent RNA chain at the catalytic site of RNA polymerase. Other mutations like A490E [(30)] and Δ490-495 [(31)] in RpoC have been implicated to alter NTP sensing, uptake and finally delay stable elongation complex(SEC) formation (*Supplementary Figure S10*). This would lead to a slow transcription by the mutated polymerase, and explain the slow growth exhibited by *RpoC^A494V^* in different *rpoS* backgrounds. Further, this slow transcription could internally trigger stress response, initiating the upregulation of *σ^S^* targets that we observe in ZK819 and SC backgrounds. Bacterial sigma factors are known to interact with the *β* and *β’* subunits of RNA polymerase [(32)] and recently, it has been shown that *σ^S^* directly interacts with β’ subunit of RNA polymerase, diminishing the pore size of the secondary channel [(32)]. This interaction could be affected by different allelic variants of *rpoS* (*rpoS819* and *rpoS92*).

In recent literature, there has been substantial focus on SC of pathogenic origin. Different strains of *Staphylococcus aureus* [(33)]*, Pseudomonas aeruginosa* [(34)]*, Burkholderia cepacia*[(35)]*, E.coli* [(36)], have been described to form small colony variants, some of which are genetically stable. Amongst the genetically stable variants of SC, most are metabolic mutants, defective either in electron-transport chain, or in thymidine biosynthesis pathway [(14)]. Early studies have found non-pathogenic *E.coli* to unstably form small colonies under prolonged exposure to chemicals like phenol or anthraquinone [(37, 38)]. Under our experimental framework, we observed that *E. coli* diversifies into stable, regulatory mutants which phenotypically produces small colonies with enhanced stress tolerance after three weeks of incubation in rich spent media.

In summary, we show that a mutation in the core RNA polymerase provides a competitive advantage to its bearer in deep stationary phase, by altering the balance between growth and stress tolerance.

## Materials and Methods

### Bacterial strains and culture conditions

*Escherichia coli* K-12 str W3110 is the parent strain of *Escherichia coli* ZK126. The latter, when evolved in prolonged stationary phase for 10 days gave rise to *Escherichia coli* ZK819 which has an elongated and attenuated *rpoS* allele [(4)] called *rpoS819*. ZK819 was evolved in prolonged stationary phase for 28 days [(5)]. The small colony form isolated is referred to as SC and its co-existing ancestral colony sized variant LC.

Additional long-term stationary phase experiments were performed to test repeatability of the appearance of small colony forms. Every day, 1 ml aliquots were taken from each flask. 500ul of this sample was used to prepare dilutions in 0.9% NaCl solution which were then plated on LB plates in two technical replicates. The remaining 500ul solution was mixed with equal volume of 50% glycerol and saved in −80°C as stocks. (Figure 1). The plates were incubated at 37^0^C for 16-18 hours and the colonies were counted to track population changes over time and to check for contamination. Images were taken with the help of Gel-Doc system. Putative small colonies, whenever seen, were re-streaked to verify stability of the phenotype and a separate count of them was maintained daily.

For measuring growth characteristics of various strains, 5 ml of LB was inoculated with single colonies of respective strains and grown overnight to form the starter culture. This culture was diluted 1000 fold in 30 ml LB in 100 ml conical flasks. Cultures were maintained at 37^0^C, with circular shaking at 200 rpm to maintain aeration. Growth dynamics were tested in rich media (Luria broth, Himedia), as well as minimal media (M9) with 0.5% glucose as carbon source. Bacterial growth was measured by tracking change in optical density at 600 nm.

### Calculation of maximum growth rate and lag time

To estimate the maximum growth rate, specific growth rates of each strain was calculated from turbidimetric change for every 15 minutes over 20 hours. Based on the extent of initial noise a sliding window of size 5-8 timepoints was taken for each strain and the median growth rate was ascertained for each window. The element of the window showing maximum median was defined as corresponding to the maximum growth rate for a specific strain.

To calculate time spent by each culture in lag phase the last point of lag was determined following Bertrand (2019) [(39)]. Intersection of non-growing lag phase and exponentially growing log phase was ascertained by fitting linear models to growth dynamicss. The time obtained from this intersecting point was considered as the duration of the lag phase for a specific strain.

**Table1:** Strains used in this study

| Strain | Description | Source |
| --- | --- | --- |
| *Escherichia coli* W3110 | Parent | CGSC #4474 |
| ZK126 | W3110 lac U169 *tna-2* | [5] |
| ZK819 | ZK126 *rpoS819* | [5] |
| Large Colony Variant(LCV) | Evolved strain, *rpoS<sup>WT</sup>*, *icd<sup>H366H</sup>* | This Study |
| Small Colony Variant(SCV) | Evolved strain, Small colony size,<br>*rpoS+92*, *rpoC<sup>A494V</sup>* | This study |

### Construction of strains with rpoC^A494V^ mutation

Transferring and reverting the point mutation *rpoC^A494V^* in different genetic backgrounds was performed using *Escherichia coli* str CAG18500 (CGSC; *thiC39*) [(18)]. In this strain, *thiC,* a gene 4.6kb upstream of *rpoC,* is tagged with tn10 carrying a tetracycline resistant marker. Co-transduction frequency of *rpoC* in this strain has been previously reported as 65% [(18)]. P1 transduction, using this strain as donor, was used to first transfer the selection marker to *rpoC^A494V^* - carrying strain. Among transductants having the selection marker, one set carried *rpoC^A494V^* and the rest *rpoC^WT^*. The former were used as donors for further P1 transduction of the mutation into different genetic backgrounds. The transductants were plated on LB agar having 12.5mg/ml of tetracycline and were screened for the presence or reversion of the point mutation by Sanger sequencing [primer sequences supplied in *Supplementary Table S2*]. Marked strains carrying wild type *rpoC* gene was utilized as donor to revert the mutant allele where applicable.

### Determination of Minimum Inhibitory Concentration(MIC) of SC and ancestor strains against different antibiotics

100μg/ml stock solutions of relevant antibiotics were prepared and serial dilutions were made for each antibiotic within a range of 0 to 100 μg/ml in 300μl of LB. 200μl LB with the appropriate antibiotics was dispensed into wells of a 96-well plate, which was then inoculated with overnight grown cultures of appropriate strains in 1:100 dilution. The plates were incubated for 24 hours at 37^0^C, 200 rpm. Turbidity of the cultures were checked by a spectrophotometer at 600nm. OD at 600nm was plotted as a function of antibiotic concentration and the lowest concentration of the antibiotic which produced no perceptible growth at a threshold of O.D 0.05 was taken as Minimal Inhibitory Concentration for that antibiotic. Experiment was repeated over 6 independent replicates for each strain.

Additionally, LB-agar plates containing a specific antibiotic were prepared at concentrations ranging from 0 to 100 μg/ml. Overnight grown cultures were diluted 6 to 8 folds in saline solution and plated on LB plates with different antibiotic concentration. After incubation at 37^0^C for 16 to 18 hours, the resulting colonies were counted. From each plate, the number of colony forming units was determined per unit volume and the same was plotted as a function of antibiotic concentrations. The lowest concentration of antibiotic which did not exhibit any colony growth was taken as MIC for that antibiotic for a specific strain.

### Competition experiment in fresh and spent media

Mixed culture competition experiments [(4)] were carried out to test the fitness advantage of the mutant strain compared to wild type strains in different backgrounds. Competing strains were tagged with resistance markers for kanamycin or tetracycline in a featureless genomic region between *htrC* and *rpoC*. They were then inoculated in 5ml of LB without antibiotics in 50ml falcons. Competing strains were inoculated in either equal frequencies (500 microlitres of each in 5ml LB), or one of them in 1000-fold excess (1 ml and 1 ul in 5ml media). As a control, they were inoculated in isolation to probe population dynamics under no competition. These falcons were then maintained for 5 days in standard growth conditions. Every day, cultures were appropriately diluted in saline and plated on kanamycin (50 mg/ml) and tetracycline (12.5mg/ml) supplemented LB agar plates. Colonies developed after overnight incubation at 37^0^C, were counted.

In addition to freshly autoclaved media, competition experiments were carried out in media spent by the ancestor ZK819. 3, 10, 20 and 30 day old media were used for competitions for all backgrounds. To prepare spent media, 500 ml of LB in a 1 l flask was inoculated with overnight grown culture of ZK819 in 1:100 dilution. This flask was incubated in 37^0^C, with circular shaking at 200 rpm. After appropriate growth, 100 ml of the culture was taken from the flask. Bacterial cells were pelleted by spinning the culture at 4000 rpm for 15 minutes. The supernatant was then passed through a 0.45 micron membrane filter (Sartorius) and stored in sterile 500 ml bottles.

### Arabinose induction and estimation of araB mRNA by qPCR

Appropriate cultures were incubated overnight in rich media and were diluted 100 folds in 100ml of LB. They were allowed to grow till mid-exponential phase. Induction was performed at maximum growth rate stage with L-arabinose (Himedia) added to a final concentration of 100mM in the culture. Transcription “Stop” solution was prepared following Vogel et al and pre-chilled in a −30^0^C bath [(40)]. Post induction, 500ul samples were periodically collected and mixed with equal volumes of stop solution in a −30^0^C bath. After 30 minutes incubation whole RNA fraction was isolated from each samples by Trizol (Ambion) based extraction.

qPCR was performed to amplify specific region of *araB* and *rpoA* transcripts with relevant primers (*Supplementary table S1*). One step TB green Primescript RT-PCR kit II (Takara, RR086B) was used for qPCR reaction on Quantstudio 6 Flex system (Applied Biosystems). 50μg of total RNA was used for each reaction. *araB* transcripts were estimated and internally normalized by *rpoA* transcripts.

### Whole genome and whole transcriptome sequencing

Whole genomes of SC, LC and ZK819 were isolated from single colonies, in replicates and grown overnight in LB. Whole genome isolation was performed using GenElute bacterial DNA isolation kit (Sigma-Aldrich). Paired-end sequencing was done by using the Illumina Miseq platform; libraries for the same were prepared with Truseq DNA Nano library preparation kit (Illumina) as suggested by manufacturers.

Sequence data were mapped to reference sequence of *Escherichia coli* K-12 str W3110 (GenBank accession no. NC007779.1). The Breseq pipeline ((41); www.barrciklab.com/Breseq), which uses Bowtie for read alignment, was used to call SNPs, indels and structural variants [33]. Mutations present in only SC at 100% frequency and not present in either LC or ZK819 were considered as unique to SC.

To verify whether *rpoC^A494V^* transductant strains harboured only the said mutation as compared to donors, whole genomes of transductants from five different *rpoS* backgrounds were sequenced. *rpoC^WT^* and *rpoC^A494V^* strains in *ZK126*, *ZK819*, *LC*, *SC* and *ΔrpoS* backgrounds were sequenced in duplicates in two different stages of growth. Cells were isolated during maximum growth rate which occurs just before the population enters exponential growth, and during the stationary phase of growth.

To look at the effects of the *rpoC^A494V^* mutation on gene expression, whole RNA fractions were isolated from the cell lysate of wild type and *rpoC^A494V^*-transductants in different backgrounds. Whole-RNA isolation was performed after the bacterial cultures entered stationary phase, upon 16 hours of growth. RNA isolation was performed using Trizol reagent (Ambion). mRNA fraction from whole RNA was enriched using MicrobeExpress kit (Ambion). Purity of whole RNA was checked by assessing A260/A280 ratio in Nanodrop and further on Bioanalyser. rRNA contamination was checked on 1% agarose gel and on Bioanalyser. Sequencing was carried out in Illumina HiSeq2500 platform and library preparation was done using TruSeq Stranded mRNA library preparation kit (Illumina).

RNA-seq data were aligned to the genome of *Escherichia coli* K-12 str W3110 (GenBank accession no. NC007779.1) using BWA (42). Read coverage per gene was calculated using SAMTOOLS (v.1.2) and BEDTOOLS (v2.25.0). EdgeR pipeline (Bioconductor), was used to find differentially expressed genes in mutants(*rpoC^A494V^*) compared to wild type(*rpoC^WT^*) in different backgrounds. Normalization and calling of differentially expressed genes was done according to Smyth et al [(43)]. A p-value cutoff of 0.05 was applied to identify differentially expressed genes.

### Estimating *σ^S^* levels with Western blotting

Intracellular levels of *rpoS* protein was estimated between *rpoC^WT^* and *rpoC^A494V^* strains in ZK819 background. Whole protein was isolated from respective cultures grown overnight by Tri-chloro-acetic acid (TCA) extraction. 100μg of whole cell lysate from each strain was run on NuPage 4-12% bis-tris protein gel (Thermo Scientific) at 120V for 3hours. *rpoS* protein was probed by anti-mouse IgG anti-*E.coli* RNA sigmaS clone:1RS1 antibody (Biolegend; batch-B274469). Protein gel was cut at 50KDa ladder mark. One part was used to probe protein of interest by transferring to Nitrocellulose membrane and the rest was stained with Coomassie-blue and imaged under UV fluorescence. Horseradish peroxidase(HRP) conjugated IgG (Sigma) was used to develop the blot. HRP signal was detected using SuperSignal West Dura Extended Duration Substrate (Thermo). RpoS signal from each lane was normalized by the total protein for that lane.

## Data Availability

RNA sequencing data can be accessed from Gene Expression Omnibus under accession number <u>GSE141883</u>. Whole genome sequencing data can be accessed from NCBI Sequence Read Archive under accession number SUB6656032<u>21.</u>

## Acknowledgement

We thank Awadhesh Pandit from Genomics Facility, NCBS for helping with NGS, We thank Vineeth Vengayil for help with Western blotting experiments and Farhan Ali for help with growth rate and lag phase analysis codes. We are grateful to Dr Victoria Shingler for her valuable technical inputs for transferring rpoCA494V mutation to different genetic backgrounds and pointing towards the relevant donor strain. We thank The Coli Genetic Stock Center (CGSC) for supplying us strain #CAG18500. This work was supported by DBT grant (BT/PR3695/BRB/10/979/201) and a Wellcome Trust-DBT India Alliance Intermediate fellowship (IA/I/16/2/502711) and PN is supported by NCBS Graduate programme scholarship. SC is supported by DST-SERB young scientist grant SB/YS/LS-148/2014.

